# Long-term Reconstruction of Human Airway Epithelium-like Structure *in Vivo* with hESCs-derived organoid cells

**DOI:** 10.1101/2020.07.18.209940

**Authors:** Yong Chen, Le Han, Shanshan Zhao, Jianqi Feng, Lian Li, Zhili Rong, Ying Lin

**Author notes:** These authors contributed equally: Yong Chen, Le Han. Corresponding author: Ying Lin.

## Abstract

Human embryonic stem cells (hESCs) derived lung organoids (HLOs) provide a promising model to study human lung development and disease. However, whether HLO cells could reconstitute airway epithelial structure *in vivo* remains unclear. Here we established an orthotopic xenograft system for hESCs-derived HLOs, enabling stable reconstruction of human airway epithelium *in vivo*. Removal of the mouse airway epithelium by naphthalene (NA) treatment enabled xenografted organoid cells survival, differentiation, and reconstruction of airway pseudostratified epithelium in immune-compromised NSG mice. Compared to unsorted pool cells, CD47^high^ cells generated more ciliated cells and possessed thicker pseudostratified epithelium. RNA-seq data revealed that CD47^high^ cells highly expressed epithelial cell, lung progenitor, lung proximal cell and embryonic lung development associated genes. These data reveal that HLOs hold cell therapy potential in regenerative medicine by long-term reconstituting airway epithelium.

---

Human pluripotent stem cells (hPSCs) derived lung organoids as a promising tool for human lung development study and disease modeling, as well as cell source for lung regeneration[1-5]. Although hPSC-derived epithelial bud tip organoids can engraft into an injured mouse airway and undergo multilineage differentiation, reconstruction of sequential airways epithelium structure is absent[6]. In the present study, we long-term reconstruction of sequential airways epithelium-like structure in naphthalene (NA-induced acute lung injury mice by intratracheal transplantation of lung organoid cells derived from human embryonic stem cells (hESCs).

It was reported that damaging the lung epithelium promoted engraftment of adult human lung epithelial cells[7], and that long-term engraftment (6W) of hPSCs derived bud tip organoids cells possessed differentiation capability and generated patch structures in airways of NA-injured immunocompromised NSG mice[6]. We tried to investigated differentiation and airway epithelium structure reconstruction by using our previously reported modified hESCs-derived lung organoid (HLO) differentiation protocol[1]. To track transplanted cells, a GFP reporter H1 hESC line was used in our experiments (Fig 1A). We found that pool and CD47^high^ transplantation group possessed regional GFP expression in lung lobes of NA-injured NSG mice 5 weeks after cell transplantation (Fig 1B-D). Hematoxylin and Eosin (HE) staining and GFP fluorescence microscopy results confirmed that pool and CD47^high^ transplantation groups possessed GFP expression (Fig 1E), and GFP+ cells survived on the surface of airway and possessed ∼50% lumen area (Fig 1E&F). GFP expression was also confirmed by human nuclei antibody (HUNU) co-staining (Fig 1G). Immunostaining assay results showed that pool and CD47^high^ cells both differentiated into lung proximal cell types, including ciliated cells (acetylated tubulin+ GFP+), club cells (CC10+ GFP+) and basal cells (P63+ GFP+, only in pool group) (Fig 2A&B). Furthermore, high expression of lung progenitor marker NKX2.1 were observed in both pool and CD47^high^ cell transplanted groups (Fig 2C). Together, these results demonstrated that HLOs-derived donor cells survived and differentiated into airway epithelial cells in NA-induced acute lung injury NSG mice.

**Figure 1.**
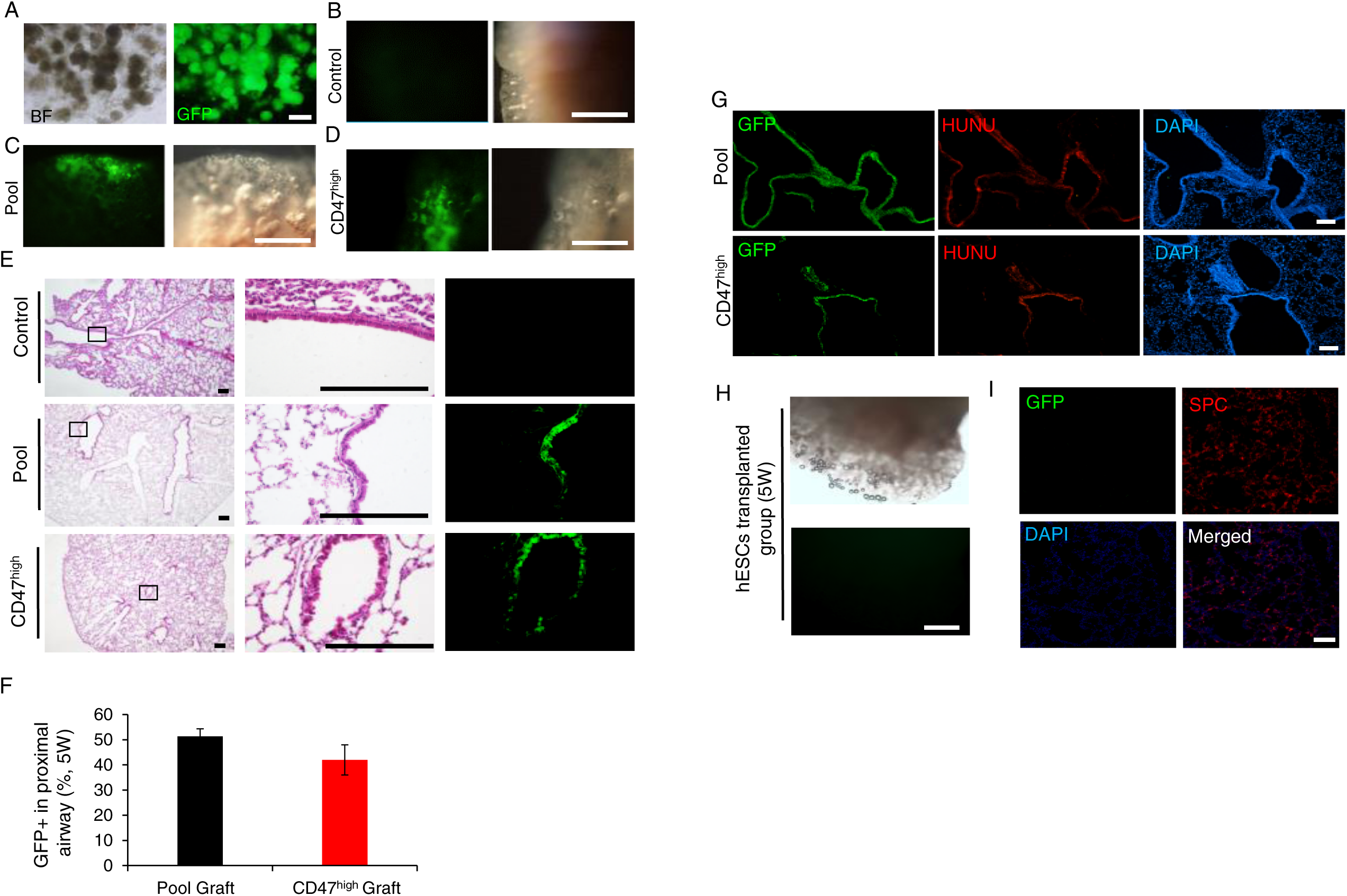
Human lung organoid-derived donor cell survival in naphthalene-induced lung injury NSG mice. (A) Morphology of GFP-labeled H1 hESCs-derived human lung organoids (HLOs). Scale bar, 100µm. (B-D) GFP expression in DPBS control group, pool group and CD47^high^ group 5 weeks after transplantation. Scale bar, 500µm (E) HE staining of lung lobe in DPBS control group, pool group and CD47^high^ group 5 weeks after cell transplantation. Scale bar, 100µm. (F) GFP+ cell ratio in lung proximal airways of transplanted NSG mice. n=5-10 sections from 3-5 mice, Data represent mean ± S.E.M. (G) Immunostaining for human nuclei (HUNU) in pool and CD47^high^ groups 5 weeks after transplantation. Scale bar, 100µm (H) GFP expression in GFP-labeled H1 hESCs transplanted group 5 weeks after transplantation. Scale bar, 500µm. (I) Immunostaining for SPC and GFP in GFP-labeled H1 hESCs transplanted group 5 weeks after transplantation. Scale bar, 20µm.

**Figure 2.**
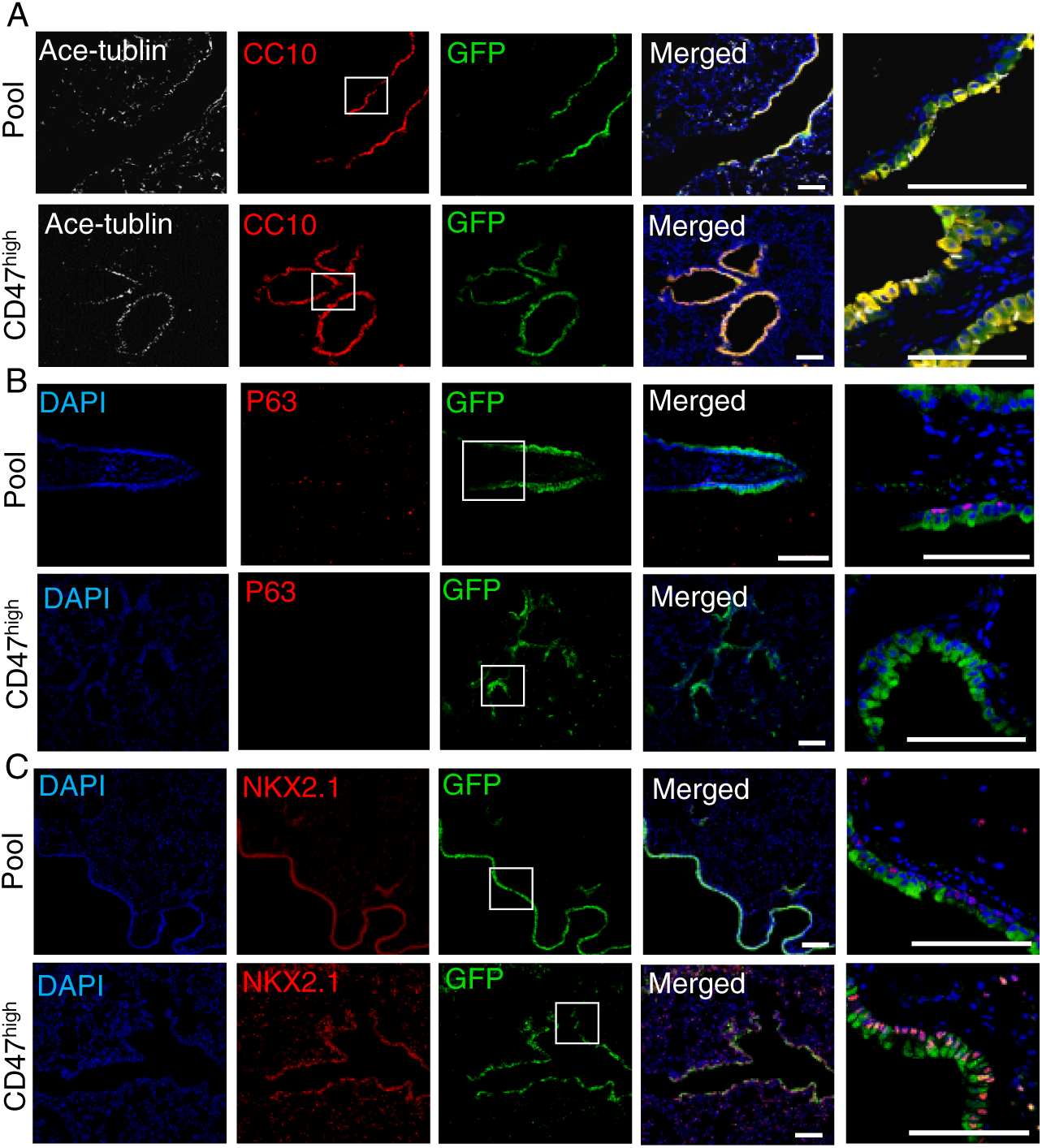
Differentiation of donor cells into proximal airway cell types. (A-C) Immunostaining for acetylated tubulin (ciliated cells marker), CC10 (club cell marker), p63 (basal cells marker), NKX2.1 (lung progenitor marker) and GPF in pool and CD47^high^ groups 5 weeks after cell transplantation. Scale bar, 20µm.

To assess donor cell survival, differentiation and reconstruction of sequential airways epithelium-like structure in NA-injured NSG mice, we examined the mice in a long-term engraftment (16 weeks) experiment. Stereoscopic fluorescence microscope analysis revealed GFP expression in lung lobes of recipient model mice (Fig 3A&B). Donor cells differentiated into ciliated and club cells and maintained sequential airway epithelium-like structure (Fig 3C). Further analysis revealed that CD47^high^ cells transplanted group generated more ciliated cells and possessed taller cell height of pseudostratified epithelium than the pool group (n=3, \**p* < 0.05, two-tailed Student’s *t* test) (Fig 3D&E). To investigate the feature of CD47^high^ cells, we performed RNA-seq analysis of CD47^high^ and CD47^low^ cells sorted from D25 HLOs. RNA-seq data showed that CD47^high^ cells highly expressed epithelia cell associated genes (*E-CADHERIN, EPCAM* and *ZO-1*), lung progenitor and proximal associated genes (*SOX2, FOXA2* and CC10) (Fig 4). These data indicated CD47^high^ cells were lung lineage determined cells.

**Figure 3.**
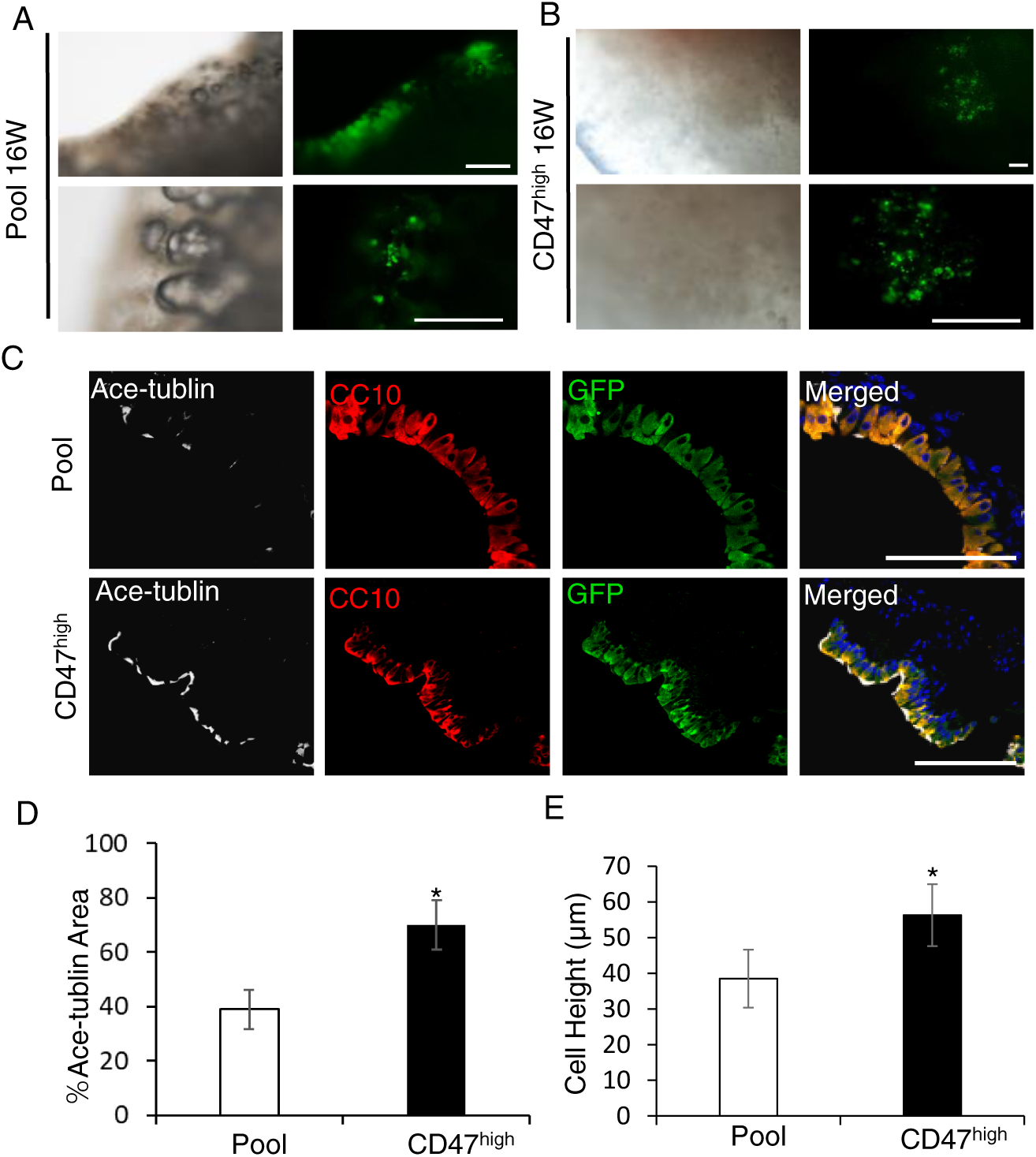
Long-term survival of donor cells and reconstruction of airway epithelium-like structure in naphthalene-injured NSG mice. (A-B) Expression of GFP in pool and CD47^high^ cell transplanted groups 16 weeks after transplantation. Scale bar, 500µm (C) Immunostaining for acetylated tubulin (ciliated cells marker), CC10 (club cell marker) and GPF. Scale bar, 20µm. (D) Ratio of donor cells derived ciliated cells in pool cells and CD47^high^ cell transplanted groups 16 weeks after transplantation, n=3, *P < 0.05, compare to Pool group. (E) Cell height of donor cells derived epithelium like structure after intratracheal transplantation for 16 weeks, n=3-5, \**p* < 0.05, two-tailed Student’s *t* test.

**Figure 4.**
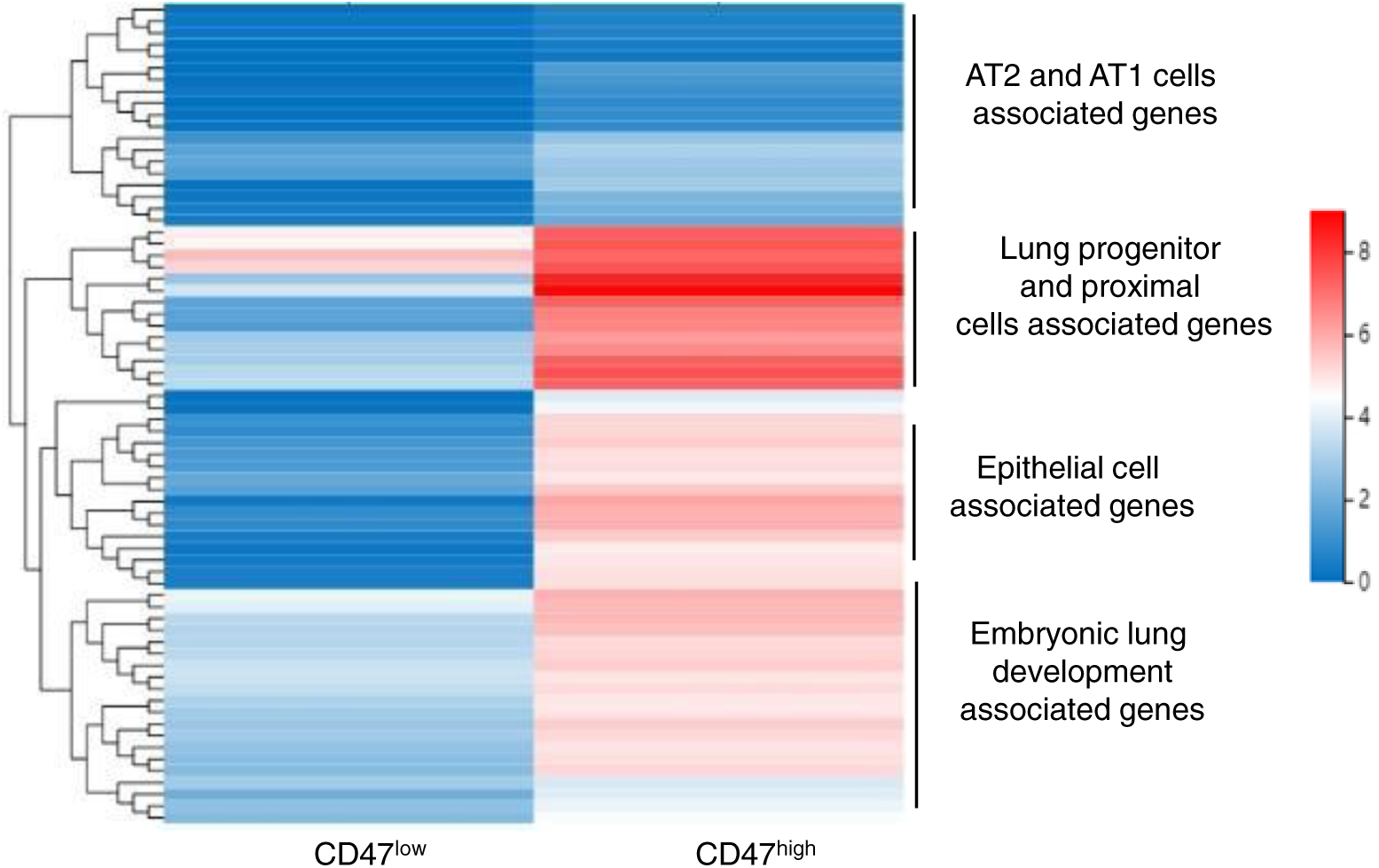
Hierarchical clustering heatmap of differential expressed genes related to lung development between HLO-derived CD47^high^ and CD47^low^ cells. hESCs were directed-differentiated to lung organoid, and HLOs at day 25 were harvested for RNA-seq analysis.

In the present study, we reconstructed human airway epithelium-like structure in vivo by using hESCs-derived lung organoid cells. Compared with the previous report that hPSC-derived epithelial bud tip organoid cells could form patch structures along the airway epithelium [6], in our study, engrafted hESCs-derived lung organoid cells could survival, differentiate and long-term reconstruct airway epithelium-like structure. It has been reported that CD47^high^ cells were NKX2.1+ cells[8], and our RNA-seq data indicated that CD47^high^ cells in D25 lung organoids were lung lineage determined cells. The strategy in the present study to reconstruct human airway epithelium structure *in vivo* provides a potential tool to investigate cell therapy and lineage tracing, as well as SARS-CoV-2 study.

## Materials and Methods

### Maintenance of hESCs

GFP labeled H1 hESCs were cultured on Matrigel (BD Biosciences, 354277) coated plates in mTeSR1 medium (StemCell Technologies, 05850) at 37°C with 5% CO_2_. Cells were manually passaged at 1:4 to 1:6 split ratios every 4 to 5 days. All hESC work in this study has been approved by the Institutional Embryonic Stem Cell Research Oversight Committee (ESCRO) of Southern Medical University.

### Generation of hESCs derived lung organoids

hESCs derived lung organoids were generated as our previously described[1]. Briefly, definitive endoderm cells were generated from hESCs (∼90% confluence) by the combination of 1-2µM CHIR-99021 (Tocris, 4423-10MG) [9] and 100ng/ml ActivinA (R&D Systems, 338-AC-050) for 3 days. Then 200ng/ml Noggin (R&D Systems, 6057-NG-100), 500ng/ml FGF4 (Peprotech, 100-31-1MG), 10µM SB431542 (Tocris, 1614-10MG) and 2µM CHIR99021 (Tocris, 4423-10MG) for 4 days [3]. After 7 days treatment with factors, cells were embedded in a droplet of Matrigel (BD Biosciences, 356237) and fed with Advanced DMEM/F12 containing 1% fetal bovine serum. Organoids were transferred into new Matrigel droplets every 5-8 days.

### Flow cytometry

The cells (D25, lung lineage organoids) were incubated in Accutase for 20 min at 37°C, then collected gently. For CD47^high^ cells sorting, cells were blocking with 10% FBS solution at RT for 1 hour. Primary antibodies were added at appropriate dilutions for 30 min at RT. After rinsing with 2% FBS/PBS, the cells were analyzed using a BD LSRFortessa flow cytometer (BD Biosceiences).

### Naphthalene-induced lung injury mice model and cell transplantation

Immune deficient B-NSG (NOD-Prkdcscid IL2rgtm1/Bcgen) mice were purchased from Beijing Biocytogen Co., Ltd. (Beijing, China). Naphthalene (NA) (200mg/ml) was intraperitoneal injected once as previous described [6]. Intratracheal transplantation of D25 HLOs derived pool and CD47^high^ cells (2-5×10^5^ per model mice). Lung tissues were harvested 5 and 16 weeks after cell transplantation. There were normal group, DPBS control group (model mice), hESCs transplanted control group, pool cells transplanted group and CD47^high^ cells transplanted group, n=8-12 for each group.

### Immunofluorescence staining

Samples were fixed in 4% PFA overnight at 4°C, then rinsed with PBS and incubated overnight again at 4°C in 30% sucrose solution. The samples were next overlaid with OCT compound and frozen using dry ice and stored at -80°C. 10μm sections were permeabilized with 0.2% Triton X-100 (Sigma, T9284)/PBS for 45 min at RT, and then blocked with blocking solution (10% FBS) at RT for 1 hour at 4°C. Primary antibodies were added at appropriate dilutions overnight at 4°C, and secondary antibodies at RT for 1 hour. Finally, samples were counterstained with DAPI (Sigma, D9542) for 5 min, and then imaged with the Zeiss LSM 880 confocal microscope (Carl Zeiss). Antibodies used in this study are listed in Table S1.

### Hematoxylin and Eosin staining

10μm Paraffin section of lung tissues were used to perform Hematoxylin and Eosin (HE) staining following the instruction, and then imaged with Nikon microscope.

### Experimental replicates and statistics

All experiments were done for at least three (n=3) independent biological replicates. Statistical analysis were done using Prism software. If only two groups were being compared, a two-tailed Student’s *t* test was employed. The differences between the groups were considered statistically significant for *p* ≤ 0.05.

## Acknowledgments

This research was funded by the National Natural Science Foundation of China (81872511 and 81670093), Frontier Research Program of Guangzhou Regenerative Medicine and Health Guangdong Laboratory (2018GZR110105005), the Program of Department of Science and Technology of Guangdong Province (2014B020212018), the Natural Science Foundation of Guangdong Province (2018A030313455).

## Figure Legends

**Table S1.** Antibodies used in the study.

| ***Antibodies*** | ***Dilution rate*** | ***Manufacturer*** | ***Cat. No.*** |
| --- | --- | --- | --- |
| NKX2.1 | 1:250 | Abcam | ab76013 |
| HUNU | 1:400 | EMD-Millipore | MAB1281 |
| P63 | 1:200 | Abcam | ab124762 |
| CC10 | 1:300 | Abcam | Ab40873 |
| Acetylated TUBULIN | 1/1000 | Sigma | T7451 |
| Mature-SFTPC | 1/800 | Seven Hills | WRAB-76694 |
| Donkey anti-goat ( RRX ) | 1:500 | Jackson ImmunoResearch | 705-295-147 |
| Donkey anti-goat (Alexa488) | 1:500 | Jackson ImmunoResearch | 705-545-147 |
| Donkey anti-rabbit (Alexa488) | 1:500 | Thermo Fisher Scientific | A-21206 |
| Donkey anti-mouse (Alexa647) | 1:300 | Thermo Fisher Scientific | A-31571 |
| Donkey anti-mouse (Alexa568) | 1:500 | Thermo Fisher Scientific | A10037 |

## Notes

### Competing Interest Statement

The authors have declared no competing interest.

